# Modular Parallel Plate Flow Chamber with Tunable Substrate Mechanics and Defined Shear Stress

**DOI:** 10.1101/2025.11.26.690741

**Authors:** Bryan J. Ferrick, Jason P. Gleghorn

## Abstract

Cells integrate multiple mechanical cues simultaneously, yet most *in vitro* models examine extracellular matrix (ECM) stiffness and fluid shear stress (FSS) in isolation, limiting our understanding of mechanotransduction. We developed a parallel plate flow chamber with a polyacrylamide (PAA) substratum enabling independent, tunable control of substrate stiffness and FSS using readily available materials. We confirm that the PAA substratum has controllable mechanical properties that support the growth of Madin-Darby canine kidney epithelial cells across a range of stiffnesses. Furthermore, the flow chamber design accommodates the volumetric equilibrium swelling of the gel, maintaining a predictable fluid channel height that allows for the application of controlled fluid shear stress to cells within the device, confirmed through particle image velocimetry of perfused microspheres. Single flow chambers support the growth of sufficient cellular numbers for endpoint analyses, such as Western blots. Finally, quantitative analysis of F-actin organization revealed that substrate stiffness and FSS synergistically increase filament length with independent effects on filament width, demonstrating the ability and usefulness of this model as a tool for studying the effect of multiple concurrent forces on cell behavior.

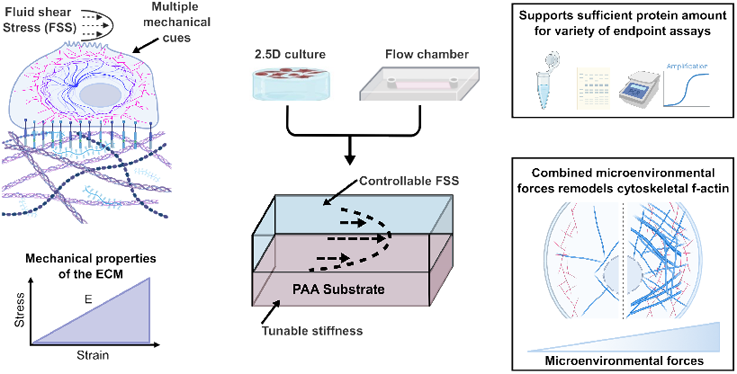

## Introduction

Mechanical forces within the tissue microenvironment play a central role in regulating biological functions ranging from embryonic development and tissue morphogenesis to wound healing and disease progression (1–7),. These forces, arising from extracellular matrix (ECM) stiffness, fluid shear stress (FSS), tensile strain from tissue stretch, and morphological remodeling, are sensed and integrated by cells through focal adhesions, intercellular junctions, cilia, mechanosensitive ion channels, and the cytoskeleton to control gene expression and cell function. Among these forces, ECM stiffness and FSS are two of the most widely studied and biologically potent, often acting through shared mechanosensitive pathways (8–16). For example, the Hippo pathway transcription factors Yes-associated protein (YAP) and PDZ-binding motif (TAZ) respond to both substratum compliance and local cellular density to control cell proliferation and inhibit apoptosis (17, 18), while the mechanosensitive ion channel Piezo1 is activated by both substratum stiffness and fluid drag-induced membrane tension (19).

Despite being commonly studied in isolation, these forces often act simultaneously and synergistically *in vivo*. In the renal and vascular systems, for instance, FSS and tissue compliance together regulate calcium influx and activation of signaling pathways that influence epithelial transport and vascular tone (19–22). The shear stress generated from fluid flow is crucial to kidney function, activating the water transporter protein aquaporin-2 (23), and increasing tight junction formation of mature proximal tubule cells (24). Studies have shown that Piezo1 responds to both membrane tension from substrate stiffness and deformation from flow, thus serving as a critical integrator of microenvironmental forces and influencing cytoskeletal organization (19, 25, 26). Understanding how cells integrate these concurrent mechanical cues is central to deciphering mechanisms of tissue function in both health and disease states, such as fibrosis, in which progressive ECM stiffening and altered hemodynamics occur concurrently.

Whereas animal models recapitulate the complete complement of forces acting on the tissue, precisely measuring, controlling, and modulating the biophysical properties *in vivo* is difficult. *In vitro* models can control variables more tightly, and there are many models in which ECM stiffness or FSS can be studied independently. Investigations into the effects of ECM stiffness on cell behavior and function frequently use polyacrylamide (PAA) hydrogels (27–32). PAA hydrogels can be readily prepared to control the material’s elastic modulus, making them ideal substrates for cell culture when material stiffness is a variable of interest. The addition of fluid flow in *in vitro* studies is technically challenging but can be implemented most simply using a parallel-plate flow chamber or a cone-and-plate device. In these studies, FSS is applied to cells cultured on a glass, plastic, or polycarbonate membrane substrata, and outcomes such as cell morphology or gene expression can be compared to cells not exposed to FSS. However, these substrates have mechanical properties that are orders of magnitude higher than the native *in vivo* microenvironment (19, 23, 33–39). This mismatch limits the ability to study mechanosensitive pathways that depend on physiologic stiffness. A major technical challenge is that most parallel-plate flow chambers lack physiologically relevant mechanical properties and do not permit independent, tunable control of both ECM stiffness and FSS within a single, accessible platform (40).

Although more advanced 3D microfluidic models are capable of applying controlled ECM stiffness and FSS simultaneously, they typically have lower throughput, have constrained available endpoint applications due to low cell numbers, and are often inaccessible to non-engineers due to their complexity, specialized fabrication requirements, and cost (41, 42). Additionally, some capable models require the use of vacuum pumps to seal the device for perfusion, which adds complexity (43, 44). Therefore, there is a need for a simple, accessible, and high-throughput device that allows for independent control of both ECM stiffness and FSS, enabling mechanistic studies by field experts across biology and engineering.

To address these challenges, we have developed a highthroughput parallel plate flow chamber with a polyacrylamide hydrogel substrate, enabling independent control of substratum stiffness and fluid shear stress. The device is intentionally designed to be fabricated using commonly available materials and equipment, making it accessible to a wider range of users. We validated the independent control of substrate stiffness and fluid shear stress through the rheological analysis of PAA hydrogels and particle image velocimetry of fluorescent microspheres perfused through the flow chamber, both with and without a monolayer of epithelial cells. Additionally, by measuring total protein content, a single flow chamber supports the growth of sufficient cellular material for endpoint analyses such as Western blot. Finally, we demonstrate a combinatorial effect of hydrogel stiffness and FSS on F-actin cytoskeletal remodeling in Madin–Darby canine kidney epithelial cells, highlighting the potential of the device for studies that require combined mechanical inputs.

## Materials and Methods

### Device fabrication

The flow chamber was composed of a PAA hydrogel for cell culture bonded to an amino-silanized glass coverslip (24 × 60 mm) with stacked layers of a silicone sheet (1/32 in, McMaster-Carr, 86465K21). An acrylic top cover completed the assembly, bonded together with two layers of a pressure-sensitive adhesive (PSA) tape (3M 9500PC, St. Paul, MN) to create the channel for flow (**Figure 1A**). Component geometries were designed in SolidWorks 2021 (Dassault Systèmes) and cut using a vinyl or a laser cutter to create an internal channel measuring approximately 10 × 40 mm. The acrylic top covers were cut to match the 24 × 60 mm coverslip base with two 3 mm holes for the fluidic ports positioned 40 mm apart at either end of the interior flow chamber. The fluid port holes were tapped to accept threaded Luer fittings (McMaster-Carr, 51525K235).

**Fig. 1.**
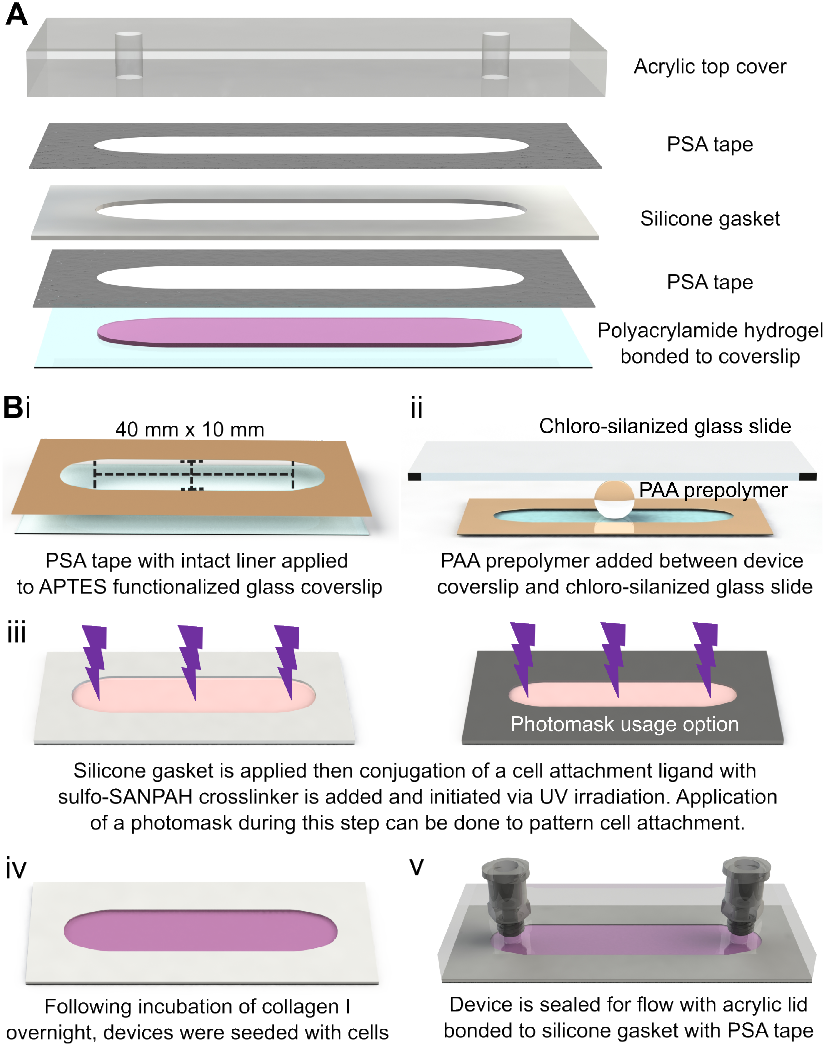
Overview and fabrication of the parallel plate flow chamber with polyacrylamide substratum. **(A)** Exploded view of the components, including the PAA hydrogel, glass coverslip, PSA tape layers, silicone gasket, and acrylic top cover. **(B)** Fabrication workflow. (i) PSA tape layer on the glass coverslip defines the hydrogel region. (ii) PAA hydrogel is polymerized between the amino-silanized device coverslip and a hydrophobic chloro-silanized glass slide. (iii) Sulfo-SANPAH activation and ECM protein conjugation enable cell attachment, with optional photomask use for patterned adhesion. (iv) Silicone gasket that is bonded to the first PSA tape layer to accommodate PAA hydrogel swelling, and cells are seeded and cultured before sealing. (v) If FSS is desired, the acrylic lid is bonded to the gasket with PSA tape to create a closed perfusion chamber.

To adhere the PAA hydrogel to the glass coverslip, the glass surface was amino-silanized. Briefly, the coverslips were plasma cleaned (1 minute on high, Harrick) to introduce hydroxyl groups, then treated with 2% (3-aminopropyl) triethoxysilane (APTES, Sigma Aldrich) in isopropanol, followed by 0.5% glutaraldehyde (Fisher Scientific) in DI water. The PAA prepolymer was prepared using the ratios from previously published protocols to achieve the desired stiffnesses (**Table 1**) (45). The prepared prepolymers were degassed for at least 30 minutes prior to polymerization.

**Table 1.** Formulations used to generate PAA hydrogels with target elastic moduli.

| Expected Stiffness (kPa) | AA (vol. %) | Bis-AA (vol. %) | 40% AA volume (μL) | 2% Bis-AA volume (μL) | DPBS (μL) |
| --- | --- | --- | --- | --- | --- |
| 0.5 | 3 | 0.06 | 75 | 30 | 895 |
| 2 | 4 | 0.1 | 100 | 50 | 850 |
| 4 | 5 | 0.15 | 125 | 75 | 800 |
| 10 | 10 | 0.1 | 250 | 50 | 700 |

The PAA gel was patterned on the amino-silanized coverslip by applying a layer of PSA tape with the flow chamber cut out and with intact backing liner to the functionalized coverslip (**Figure 1**Bi). Next, 275 *µ*L of the PAA gel prepolymer was added into the void space of the PSA tape on the functionalized coverslip under N_2_ gas and a hydrophobic, chloro-silanized glass slide was placed on top of the PSA tape to create a uniform top surface of the PAA hydrogel (**Figure 1**Bii) (45). After polymerization, the chloro-silanized glass slide was carefully removed, and the PAA hydrogel-coverslip devices were rinsed in PBS to remove any unpolymerized polyacrylamide. The silicone gasket layer was then affixed to the device after removing the PSA tape backing liner.

To functionalize the PAA gels for cell adhesion, 1 mg/mL sulfo-SANPAH in HEPES buffer (50mM pH 8.5, Sigma Aldrich) was added to the surface of the PAA gel and UV irradiated for at least 20 minutes (**Figure 1**Biii). A photomask was optionally used to spatially pattern cell attachment. For example, a photomask composed of a second silicone gasket and a PSA tape layer, shaped to cover 1 mm of the gel’s outer perimeter, was used to prevent cell attachment near the edges of the PAA gel. Following UV irradiation, gels of all flow chambers were washed in fresh HEPES to remove unreacted sulfo-SANPAH, then incubated with type I collagen (150 *µ*g/mL) overnight at 4 °C. The photomask, if used, was removed after overnight incubation. All devices were washed with DPBS to remove unbound collagen and were ready for cell seeding (**Figure 1**Biv). When application of FSS via perfusion was needed, an acrylic top cover was sealed to the silicone gasket with a PSA tape layer and connected for closed circuit perfusion via a peristaltic pump (Ismatec) at an appropriate flow rate to achieve the desired FSS (**Figure 1**Bv).

### Cellular seeding and immunofluorescent staining

Madin–Darby canine kidney (MDCK, ATCC) epithelial cells were cultured in DMEM supplemented with 10% fetal bovine serum (FBS, Cytiva) and 1% penicillin and streptomycin at 37 °C and 5% CO2. For all cellular experiments, 2.5e^5^ MDCK cells per device were seeded prior to sealing the flow chamber with the top acrylic cover. The devices were cultured under static conditions with daily media changes until confluence, approximately 2-3 days. After confluence was reached, devices were either kept as is and cultured under static conditions, or the acrylic top was added and media was perfused to induce FSS of 0.4 dyne/cm^2^ with flow rates of 0.094, 0.89, or 1.39 mL/min for glass, 0.5 kPa, or 10 kPa substrates, respectively. Both static and perfused devices were cultured for an additional 24 hrs under their respective conditions. After 24 hrs, the cells within the devices were fixed with 4% paraformaldehyde (PFA, ThermoFisher), permeabilized with 0.1% Triton-X 100 (Sigma-Aldrich), stained with Hoechst 33342 (Invitrogen), Phalloidin-iFluor 647 (Abcam), rat anti-E-cadherin (DECMA-1, Santa Cruz), and donkey anti-rat DyLight488 (ImmunoReagents).

### Rheology of PAA hydrogels

Rheological analysis of PAA samples with expected elastic moduli of 0.5 kPa, 2 kPa, 4 kPa, or 10 kPa was done to confirm the mechanical properties of the formulations used to create the hydrogel within the flow chamber device. The PAA prepolymers for each formulation were mixed, degassed, and polymerized under N_2_ gas into 20 mm diameter disks. Upon complete polymerization and equilibrium swelling in 1x DPBS, the storage and loss moduli of the gel disks were measured through constant strain (0.1%) and frequency (1 Hz) oscillation time-series testing for 10 minutes, as these strain and frequency values are within the linear region of the gel’s mechanical properties (46). The resulting storage modulus was then used to calculate the elastic modulus of the sample based on the assumption that polyacrylamide gels are purely elastic, with a Poisson’s ratio of 0.5, using the relationship between the shear modulus (G) and elastic modulus for linear elastic materials of E = 3G’ (47, 48).

### Hydrogel thickness measurements

To measure PAA hydrogel substrate thickness after overnight equilibrium swelling in DPBS, devices were fabricated as previously described, with the addition of fluorescent microspheres (0.5 *µ*m, Flash Red, Bangs Laboratories) to the hydrogel prepolymer to visualize the bulk of 0.5 kPa and 10 kPa stiffness gels. These two stiffness gels were chosen as they represent the minimum and maximum stiffnesses used herein, and the swelling is a function of crosslinking, which is proportional to stiffness and therefore should capture the range of hydrogel thicknesses observed here (46). Full-thickness z-stacks of the device and the gel were captured using a confocal microscope (5x objective, LSM800, Zeiss) at 9 locations across the device. The edge PSA tape and silicone layers served as reference layers of known thickness. The ratio between image-based measurements and the known actual thickness of the combined PSA and silicone layers was used to correct for refractive-index-induced aberrations and to calculate the thickness of the post-equilibrium swollen PAA gels. The fluidic channel height was calculated as the difference between the combined total thickness of the PSA tape and silicone gasket layers and the thickness of the PAA gel. The average hydrogel thickness of 0.5 kPa and 10 kPa PAA static devices was compared using a Student’s t-test using JMP Pro 17.

### Fluid flow analysis

As the fluid flow rate and the resultant shear stress are functions of the fluid channel height, the thickness of the 0.5 kPa hydrogel in the device was measured during perfusion at multiple flow rates corresponding to FSS of 0.5, 1, and 2 dyne/cm^2^. Measurements were taken to ensure the hydrogel thickness did not change due to the pressures associated with perfusion. Only the 0.5 kPa gel was analyzed, as it is the “softest” and would experience the effect of fluid flow at a lower pressure compared to “stiffer” PAA gels. The flow chamber devices were created with fluorescent microspheres embedded within the gel, seeded with MDCK cells, and cultured until a confluent monolayer had formed. Each device was first measured under static flow conditions, then culture media was perfused through the devices using a syringe pump (Fusion 100, Chemyx) while capturing full-thickness confocal z-stack images (5x/0.25 LSM800, Zeiss) of at least two locations in each device. The hydrogel thickness at each flow rate was compared with its static thickness using a repeated-measures ANOVA in JMP 17.

After confirming that the pressure during perfusion did not change the fluid channel geometry, we validated that the set flow rate within the flow chamber resulted in the expected fluid shear stress for the given channel dimensions by measuring the actual fluid velocity within the device over a range of flow rates (0.1 – 0.7 mL/min, 0.1 intervals). The measured fluid velocity was then used to calculate the FSS. Both 0.5 kPa and 10 kPa stiffness devices were created and cultured with MDCK cells until a confluent monolayer was formed. Using a Dragonfly spinning disk confocal microscope (Andor), high-speed time-lapse imaging captured the perfusion of fluorescent microspheres (500nm Dragon Green, Bangs Laboratories, Inc.) at approximately 150 frames per second with a finite burst protocol using a 10x/0.45 objective. Trackmate particle image velocimetry (PIV) plug-in for ImageJ was used to calculate the microsphere velocities. The measured velocities were used to calculate the shear stress for the channel geometry. Both the measured velocities and calculated shear stresses were compared to their respective theoretical values across the range of flow rates using a linear regression model in JMP Pro 17. Fluid shear stress was calculated using the parallel plate approximation: *τ* = 6*µQ*/*wh*^2^, where *τ* is shear stress, *µ* is dynamic viscosity, *Q* is volumetric flow rate, *w* is channel width, and *h* is channel height.

### Total protein quantification

The total amount of protein available for proteomic assays of MDCK monolayers grown on the PAA gel substrates of the flow chambers was quantified and compared to the amount of protein from single wells of 6- and 12-well plates. Flow chambers with 0.5 kPa substrates, 6-, and 12-well culture plates were seeded with MDCK cells and cultured in static conditions until a complete monolayer had formed within each condition, at least 3 days. Following monolayer formation, the cells were washed in DPBS, then lysed with RIPA buffer (Thermo Scientific Alfa Aesar) with 1X protease inhibitor (ThermoFisher). Each culture substrate was carefully scraped to collect all of the cell lysate, followed by centrifugation at 14,000g for 15 minutes to remove cellular debris. The total protein concentration of each sample was measured using a Pierce BCA Protein Assay (Thermo Fisher). The cell lysate samples were prepared and analyzed according to the manufacturer’s recommended pro-cedure. The protein concentration measured from the assay was then used to calculate the average total protein content within the sample volume of each cell lysate.

### F-actin quantification

To quantify the morphological changes in F-actin cytoskeleton organization associated with the combinatorial forces of FSS and substrate stiffness, MDCK cells were cultured within flow chambers with type I collagen-coated glass, 0.5 kPa, or 10 kPa PAA gel substrates and exposed to static or 0.4 dyne/cm^2^ FSS for 24 hrs after reaching a confluent monolayer. Upon completion of the culture period, the cells were immunostained for E-cadherin, F-actin, and nuclei as described above. In devices containing a hydrogel, the top cover and silicone gaskets of the flow chambers were carefully removed to allow the hydrogel to be placed cell-side down on a glass coverslip, enabling high-magnification imaging (63x/1.4 oil immersion objective, LSM800, Zeiss). Devices with cells cultured on glass substrates could be directly imaged with this objective and did not require this step. The resulting images were segmented into individual cells with ImageJ, then processed using Fsegment (49), an open-source Matlab-based program, to measure the average number and width of the phalloidin-stained F-actin filaments in each cell. Using images of individual cells and the expected actin fiber width as inputs, we used Fsegment to preprocess the images with Gaussian filtering, path opening, directional top-hat, and hysteresis thresholding to trace and segment each fiber. These image preprocessing and filament-trace algorithm parameters in Fsegment were adjusted on an individual basis to ensure that F-actin filaments were accurately traced and measured. The length of each filament and its average width, measured at 5 *µ*m intervals along the filament, were then determined. Both the average F-actin filament length and width in individual cells were compared using a 2-factor ANOVA across FSS and stiffness conditions, followed by post-hoc Tukey HSD all pairwise multiple comparisons in JMP Pro 17.

## Results

### Accessible device design supports epithelial culture on tunable substrates

To achieve wide accessibility, we designed the device using readily available, inexpensive materials and standard fabrication techniques. The modular assembly enables cells to be seeded and cultured in an open configuration before sealing with the acrylic top cover, when fluid flow is desired (**Figure 1B**), thereby reducing experimental complexity compared to sealed microfluidic systems. The use of vinyl-cut PSA tape and silicone gaskets provides flexibility in customizing channel geometry, ultimately allowing for the precise shape and size of the PAA substratum to be tailored to specific applications, thereby eliminating the need for cleanroom facilities or specialized equipment.

We evaluated the rheological mechanical properties of the PAA gels created form established formulations for 0.5 kPa, 2 kPa, 4 kPa, 10 kPa theoretical elastic moduli (E) gels to confirm their characteristics after equilibrium swelling (**Table 1, Figure2B**) (45). Additionally, PAA hydrogels require functionalization with ECM ligands via the heterobifunctional linker sulfo-SANPAH to enable cell attachment. UV activation through a photomask allows for spatial patterning of cell adhesion, which we used to prevent attachment within 1 mm of gel edges, ensuring uniform shear stress exposure by avoiding potential edge effects from fabrication defects (**Figure 2A**). The MDCK epithelial cells cultured within this device formed a consistent monolayer that remained stable in both static and perfused devices. Robust junctional Ecadherin staining was present in cultures for all the substrate stiffness and flow conditions tested (**Figure 2C**).

**Fig. 2.**
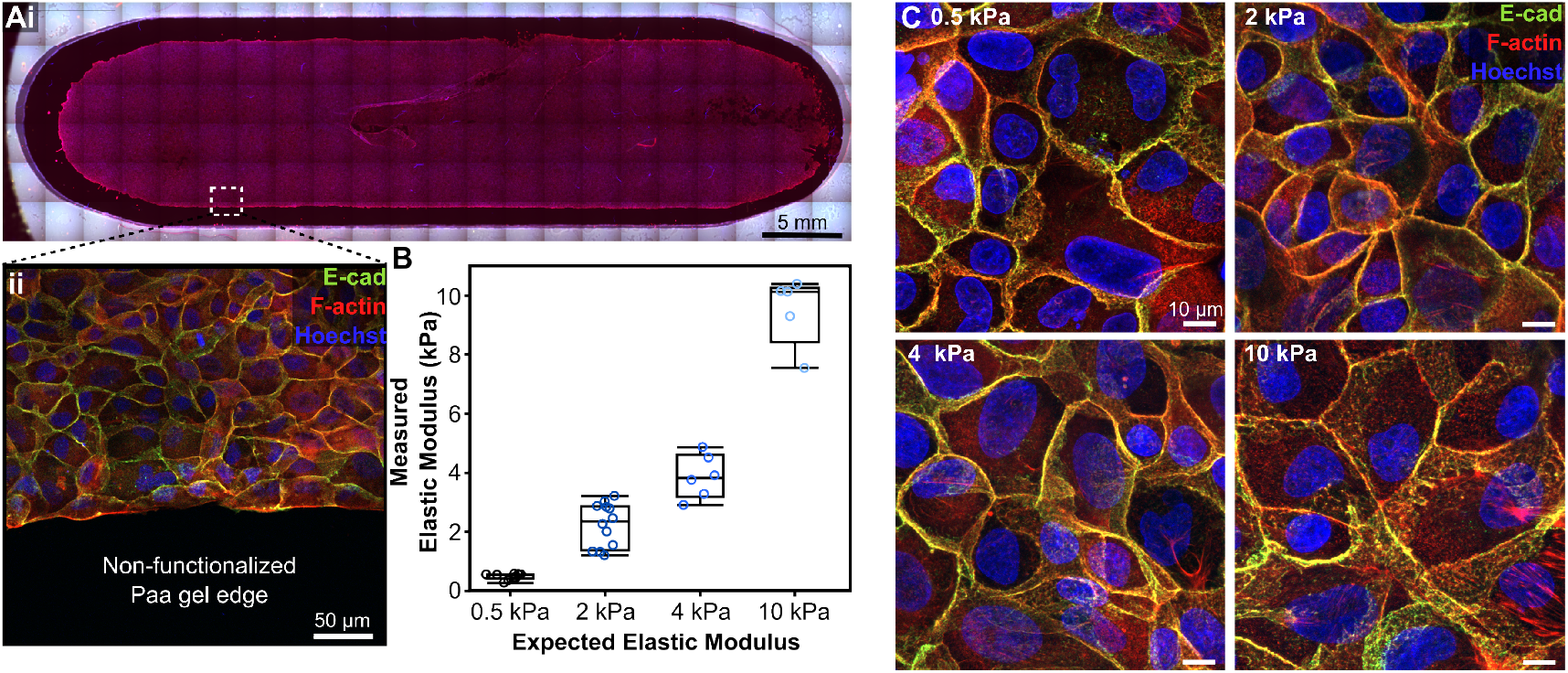
The flow chamber supports epithelial monolayer formation across PAA hydrogel substrates of varying stiffness. **(A)** Spatial ECM patterning prevents cell attachment near hydrogel edges. **(B)** Measured elastic moduli for each PAA formulation match expected modulus values (n *≥* 5 per condition). **(C)** MDCK cells form confluent monolayers with strong junctional E-cadherin staining on all stiffness substrates.

### Multi-layer design accommodates stiffness-dependent hydrogel swelling

Controlling the fluid shear stress applied to cells requires knowledge of the fluid channel geometry. This becomes challenging when the channel height is set relative to a hydrogel substrate whose thickness increases after initial polymerization due to volumetric swelling. Swelling is a common phenomenon in many hydrogel systems, including those based on polyacrylamide. To ensure the device functioned as expected, we needed to (1) accommodate swelling in the device design and (2) accurately measure post-swelling gel thickness to calculate the fluid channel height required for precise FSS application **Figure 3A**.

**Fig. 3.**
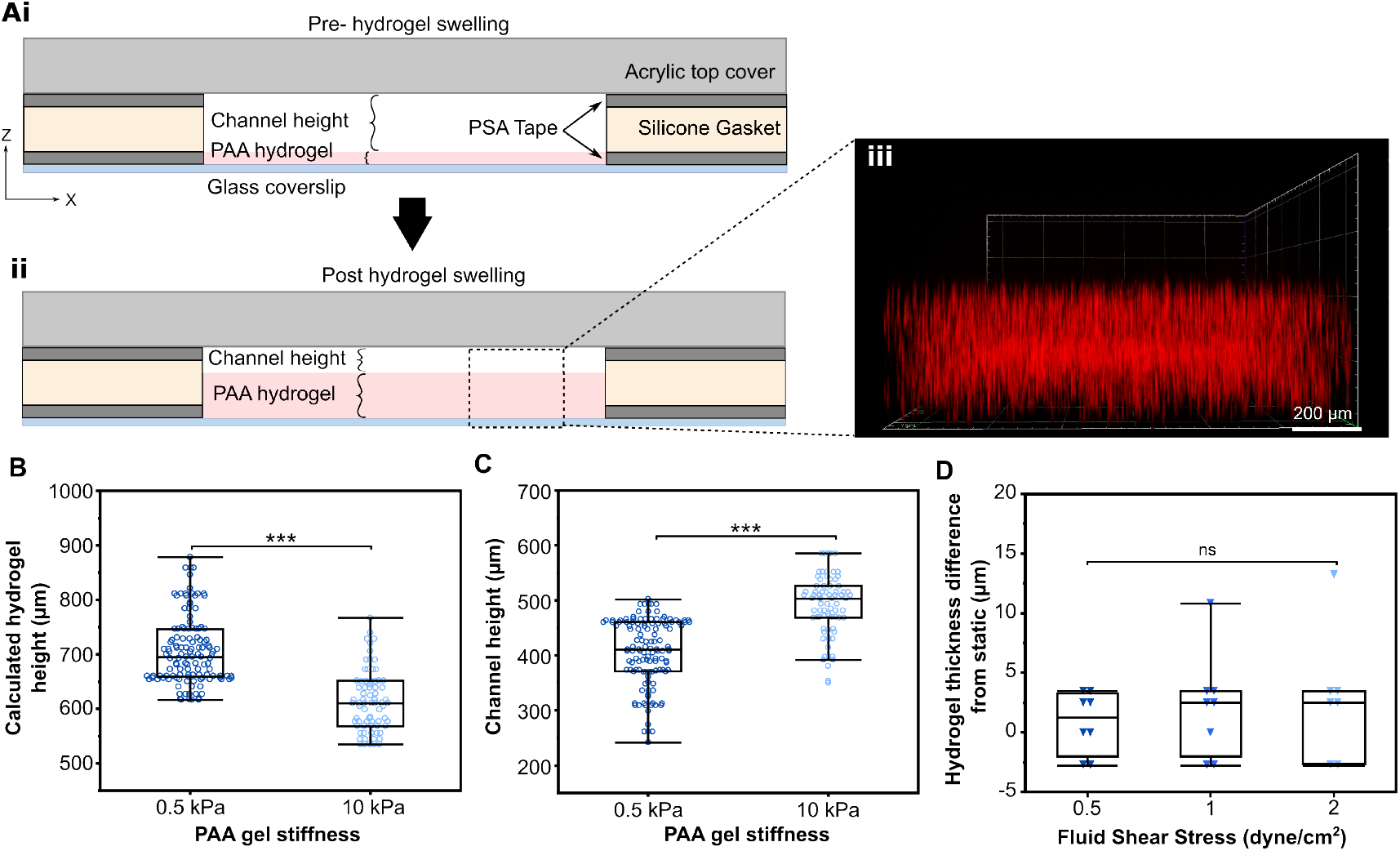
The flow chamber device accommodates the volumetric equilibrium swelling of the polyacrylamide hydrogel. **(A)** Cross-section of the flow chamber depicting the relationship between the hydrogel thickness and fluid channel height before (i) and after (ii) hydrogel swelling. (iii) ZX projection of hydrogel with embedded fluorescent microspheres used for thickness measurements. **(B)** The post swelling hydrogel thickness of 0.5 kPa gels were significantly greater than 10 kPa gels (p = 0.00011, n *≥* 7 devices, at least 7 locations per device). **(C)** The height of the 0.5 kPa PAA gels did not change significantly from their static height during perfusion (p = 0.1011, n = 4 devices, at least 2 locations per device per flow rate).

To measure the hydrogel after swelling, we captured fullthickness z-stack images of fluorescent microspheres embedded within the PAA hydrogels at multiple locations across the flow chamber devices. The ratio of known total flow chamber height to the measured thickness from the z-stack images was used to calculate the actual height of the post-swollen hydrogel. The fluid channel height was then calculated as the difference between the total gasket/tape thickness and the swollen hydrogel thickness.

Immediately after polymerization, the initial thickness of the PAA gel was 0.25 mm (250 *µ*m), as defined by the thickness of the PSA tape with its intact liner. After equilibrium swelling in PBS overnight, the mean thickness was 700.6 ± 42.8 *µ*m for 0.5 kPa gels and 600 ± 37.2 *µ*m for 10 kPa gels (**Figure 3B**), representing 2.8× and 2.4× increases, respectively. While the gels of different stiffnesses both increased in height, the amount of swelling differed. This statistically significant difference (p = 0.00011) in swelling behavior between gels of different stiffness is consistent with previous studies, as softer substrates have fewer cross-links, allowing for greater swelling (46). In our system, this swelling resulted in decreased fluid channel height for 0.5 kPa gels compared to 10 kPa gels (410.3 ± 43.2 *µ*m vs. 498.2 ± 40.4 *µ*m, p = 0.0005), reflecting an inverse relationship between gel stiffness and swelling (**Figure 3C**).

To ensure the channel height did not change due to the pressures generated within the flow chamber during perfusion, we measured the gel height of cell-seeded 0.5 kPa gel devices during perfusion across the same flow rate range. There was no significant change in gel height at any tested flow rate (p = 0.1011) (**Figure 3D**).

### Measured fluid velocities match theoretical predictions across flow rates

The magnitude of FSS applied to the cells depends on fluid velocity and, in turn, the volumetric flow rate through the channel and the channel’s dimensions. Having determined that the channel geometry remains constant during perfusion, we can validate FSS control within the flow chamber. This was achieved by measuring the fluid velocity over a range of flow rates (0.1–0.7 mL/min), which were then compared to the theoretical velocity for the corresponding set flow rate and channel dimensions of 0.5 kPa and 10 kPa stiffness devices with confluent MDCK monolayers. The fluid flow was visualized through timelapse images of 500 nm fluorescent microspheres perfused through the flow chamber, which were analyzed via PIV to determine microsphere velocities and, subsequently, the fluid velocity and fluid shear stress at each flow rate. The inclusion of the cellular monolayer was intended to mitigate fluid flow through the hydrogel bulk rather than through the channel, which could alter the velocity measurements. For both stiffness gels tested, we found close agreement between the measured and theoretical fluid velocity and calculated shear stress across the range of tested flow rates (**Figure 4**). Using high-speed imaging (150 fps), we measured fluid velocities up to 4 mm/s (0.7 mL/min), corresponding to FSS values within the physiological range for renal tubules (50). While higher flow rates have been tested within this flow chamber, accurately capturing the microsphere displacement as they convect with the fluid flow at higher velocities required a frame rate beyond the approximately 150 frames/sec available with this microscope, preventing analysis at higher flow rates.

**Fig. 4.**
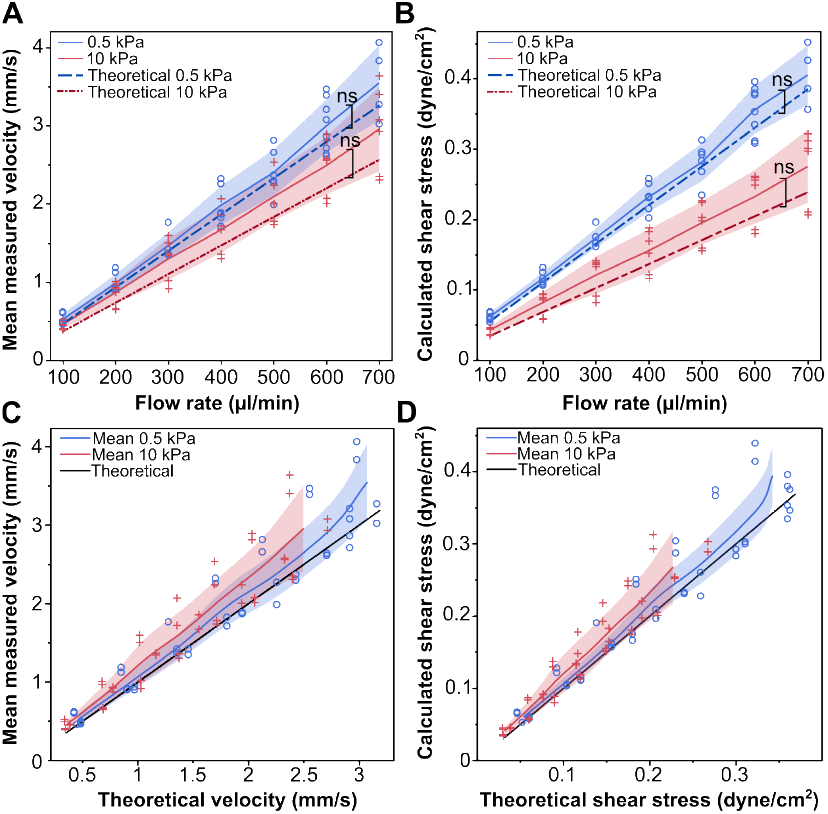
The measured fluid velocity and shear stress align with theoretical predictions for both 0.5 kPa and 10 kPa substrates. **(A)** Mean fluid velocity across tested flow rates compared with theoretical values (n = 3 devices, *≥* 2 replicates per flow rate; p = 0.222 for 0.5 kPa and p = 0.056 for 10 kPa). **(B)** Shear stress calculated from measured velocities across flow rates (p = 0.1626 for 0.5kPa and p = 0.0667 for 10 kPa). **(C)** Measured versus theoretical fluid velocity for each flow rate. **(D)** Shear stress derived from measured velocity versus theoretical shear stress. Shaded regions represent one standard deviation from the mean.

### Single channel format yields sufficient material for biochemical analyses

Many microphysiological systems generate insufficient cellular material for biochemical assays such as Western blots, limiting their utility for proteomic studies. We intentionally designed the flow chamber to provide sufficient surface area to support these assays without the need to pool samples. We cultured MDCK cells in the flow chamber and quantified the total protein content collected from the flow chamber, comparing it to the amount of protein from a well of a 6- and 1-well plate, which are more commonly used for these assays. As total protein is correlated with cell number and culture surface area, we expected the flow chamber to have less protein than a well within a 6 well plate but more protein than a well in a 12 well plate, as the culture surface areas follow that pattern (4.46 cm^2^, 9.6 cm^2^, and 3.5 cm^2^ for the flow chamber, 6- and 12-well plate, respectively) (**Figure 5A**). As expected, the total protein harvested from the flow chamber (0.32 ± 0.043 mg) was greater than that from the 12-well plate (0.17 ± 0.013 mg), but less than from a 6-well plate (0.41 ± 0.028 mg) (**Figure 5B**), reflecting the relative culture surface areas (4.5, 3.5, and 9.6 cm^2^, respectively). This amount of total protein collected from a single device is sufficient for western blot assays, which commonly require 20-40 *µ*g of protein per sample replicate (51).

**Fig. 5.**
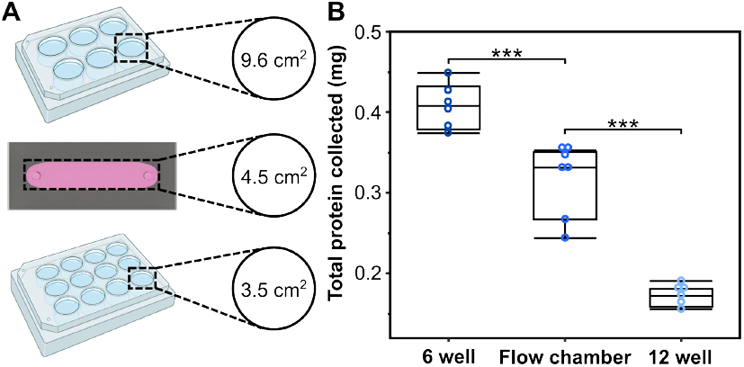
The flow chamber provides sufficient material for proteomic assays. **(A)** Comparison of available culture surface area across the flow chamber, 6-well, and 12-well formats. **(B)** Total protein yield from one flow chamber falls between that of a single 6-well and 12-well plate well (n *≥* 6 per condition; *** denotes p *≤* 0.0001).

### Substrate stiffness and fluid shear stress synergistically remodel F-actin

Remodeling of the F-actin cytoskeleton is a well-known response to changes in microenvironmental forces. This remodeling has been independently shown to occur in response to changes in ECM stiffness and fluid flow. However, the cytoskeletal remodeling in response to combined mechanical cues remains incompletely characterized. We therefore sought to quantify this remodeling in MDCK epithelial cells exposed to a combination of environmental forces by culturing them on 0.5kPa, 10 kPa, or glass substrates and simultaneously exposing them to either static or 0.4 dyne/cm^2^ FSS. The F-actin cytoskeletal organization, visualized through phalloidin immunofluorescent staining, was quantified by analyzing images of individually segmented cells with the open-source application Fsegment (49). We found an overall increase in both the average F-actin filament length and width in each cell with increasing stiffness and increasing FSS (**Figure 6**). Furthermore, the synergistic application of FSS on a stiffer substratum resulted in cells with a larger average actin filament length and width (**Figure** The increase in filament length was statistically significant for overall substrate stiffness (p < 0.0001) and FSS (p < 0.0001), as well as an interaction between the two effects(p = 0.0016). There was also a significant increase in filament width with increasing substrate stiffness (p < 0.001) and FSS (p < 0.0001), although no detectable interaction was observed between the two forces (p = 0.148). Substrate stiffness had a larger effect on filament length than FSS, while both forces had comparable effects on filament width.

**Fig. 6.**
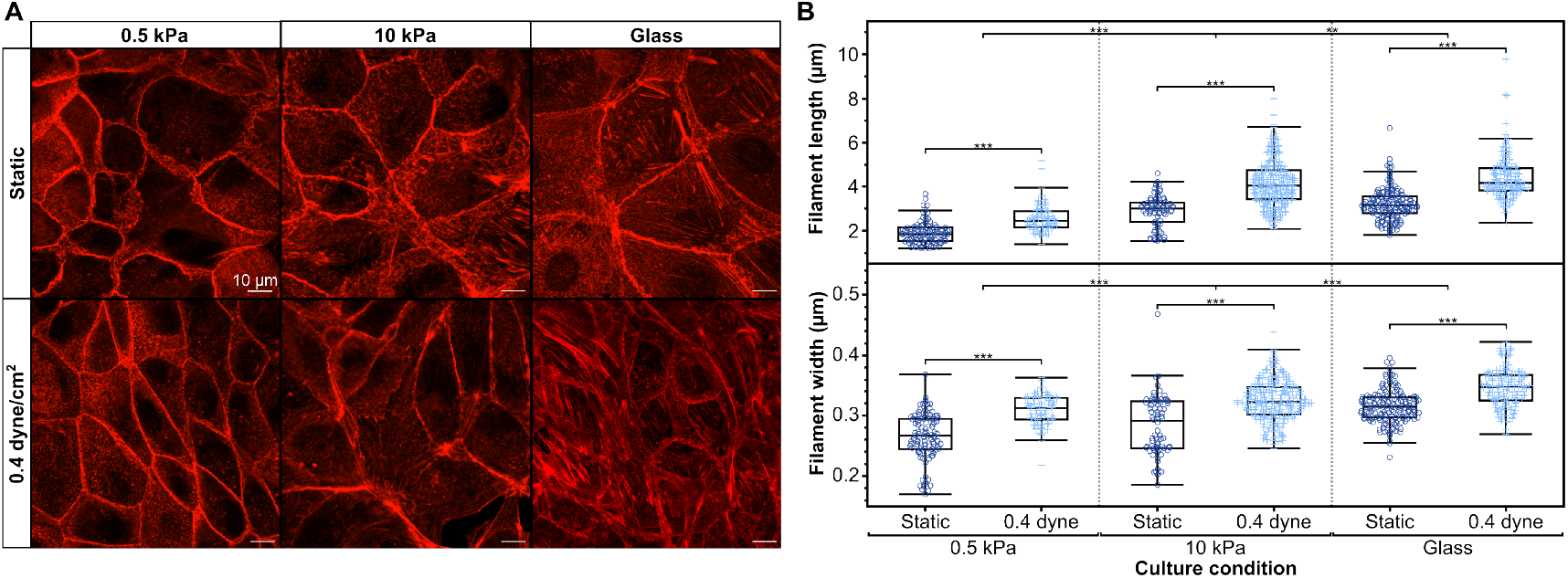
F-actin cytoskeletal remodeling in response to increasing substrate stiffness and fluid shear stress application. **(A)** Representative immunofluorescence f-actin staining of MDCK epithelial monolayers showing an increase in stress bundling of the cytoskeleton as substrate stiffness and FSS increase. **(B)** The average f-actin filament length and width within each cell of an MDCK monolayer increases as substrate stiffness and FSS increase (n *≥* 7 cells across at least 4 devices per condition; significance indicated on plots are results of post-hoc Tukey pairwise comparisons; ** denotes p < 0.001 and *** p < 0.0001).

## Discussion

F-actin cytoskeletal remodeling is a conserved response to mechanical stress that redistributes applied forces while activating mechanosensitive signaling pathways (52, 53). We demonstrate that substrate stiffness and FSS synergistically increase F-actin filament length, suggesting non-additive integration of mechanical cues. This finding is consistent with Piezo1-mediated calcium signaling, which responds to both membrane tension from substrate stiffness and fluid drag, potentially driving Rho/ROCK-dependent stress fiber formation (19, 25, 54). However, the independent effects on filament width suggest distinct downstream pathways may also be engaged, warranting further investigation of signaling networks activated by combinatorial mechanical stimuli. While we focused on cytoskeletal readouts, future studies should examine mechanosensitive signaling pathways (YAP/TAZ localization, calcium dynamics, gene expression) to provide mechanistic insights. *∼* Alternative models for studying combinatorial mechanical cues include 3D microfluidic organ-on-chip systems, which better recapitulate tissue architecture but typically yield insufficient cellular material for biochemical assays and require specialized fabrication expertise. Our flow chamber addresses these limitations by supporting protein yields comparable to 6- and 12-well plates (~320 µg per device), enabling Western blots and other proteomic analyses (**Figure 5**) (51). The use of readily available materials, simple assembly without vacuum sealing, and compatibility with standard microscopy make this platform accessible to laboratories without engineering infrastructure.

We selected polyacrylamide hydrogels for their well-characterized, reproducible mechanical properties and biological inertness. Unlike collagen gels, whose stiffness depends on temperature, pH, and source variability, PAA formulations reliably produce target moduli (55–58). The requirement for ECM functionalization, while adding a fabrication step, enables systematic investigation of specific integrin-ligand interactions (59). Photomask patterning during sulfo-SANPAH activation provides spatial control of cell adhesion, which we leveraged to prevent edge effects but could be extended to create co-culture patterns or investigate mechanotransduction at tissue boundaries. Although not tested here, the fabrication techniques can be extended to use other hydrogel systems (e.g., alginate, PEG) or cell types within the device, if desired, for tissue-specific applications. PAA hydrogels, like many other synthetic hydrogels, undergo volumetric swelling (2.4-2.8x thickness increase depending on crosslinking density) when hydrated, which complicates precise control of fluid channel geometry. The degree of swelling is linked to multiple factors, including the extent of crosslinking, mesh size, and charge within the polymeric network, which act to balance osmotic pressure and draw water into the gel (48). Our multi-layer gasket design accommodates this swelling while maintaining a predictable channel height. We developed an imaging-based measurement approach using reference layers of known thickness to correct for refractive index-induced focal shifts during confocal microscopy (60), enabling accurate determination of post-swelling gel thickness. Reference z-stack images of the tape and silicone layers with known thickness values were also captured. This was done to account for any discrepancies in reported versus actual z height due to imaging artifacts such as spherical aberrations from imaging through multiple media with different refractive indices (i.e., air, glass, and water) (61–63). This correction was essential for precise FSS application, as validated by the close agreement between theoretical and measured fluid velocities (**Figure 4**).

This platform has broad applicability for mechanobiology research. In developmental biology, it could elucidate how embryonic tissues integrate ECM remodeling and hemodynamic forces during morphogenesis. In disease contexts, the device enables modeling of pathological mechanical environments, for example, studying how fibrotic stiffening and altered blood flow synergistically promote endothelial dysfunction in atherosclerosis, or how tumor matrix stiffness and interstitial flow cooperate to drive cancer cell invasion. Whereas the current design is limited to single-channel experiments, parallelization using multi-channel devices could enable higher-throughput screening. The high throughput and compatibility with biochemical assays also make this system suitable for screening therapeutic compounds targeting mechanotransduction pathways.

## Conclusion

We developed an accessible parallel plate flow chamber enabling independent control of ECM stiffness and fluid shear stress. The device accommodates polyacrylamide hydrogel swelling while maintaining predictable fluid dynamics and protein yields sufficient for Western blots. As such, this platform overcomes a key limitation of microfluidic systems. Application to epithelial mechanobiology revealed synergistic effects of stiffness and FSS on F-actin filament length, demonstrating the utility of this approach for studying combinatorial mechanical signaling. This cost-effective, highthroughput system provides a valuable tool for mechanistic studies in development, physiology, and disease, with potential applications in drug screening and tissue engineering.

## ACKNOWLEDGEMENTS

The authors thank Katherine M. Nelson, Ph.D., for reviewing and commenting on the manuscript. This work was supported in part by grants from the National Institutes of Health: R01DE029655 and R01HL145147. Microscopy equipment was acquired with a shared instrumentation grant (S10 OD030321) and access was supported by NIH-NIGMS (P20 GM103446, P20 GM139760) and the State of Delaware. The BioRxiv template was adapted from the Henriques lab.

## DATA AND CODE AVAILABILITY

Downloadable engineering drawings for the device’s laser and vinyl cut components can be found at: https://www.gleghornlab.com/resources

## AUTHOR CONTRIBUTIONS

Conceptualization (BJF, JPG), Methodology (BJF, JPG), Investigation (BJF), Validation (BJF, JPG), Visualization (BJF, JPG), Formal Analysis (BJF, JPG), Writing – Original Draft (BJF), Writing – Review & Editing (BJF, JPG), Supervision (JPG), Project Administration (JPG), Funding acquisition (JPG)

